# Fast and slow-paced reproductive life history across native and invasive populations of a predatory marine snail

**DOI:** 10.1101/2025.05.15.654356

**Authors:** Allison L. Rugila, Emily Bucari, Emma Rawson, Amara L. Schlaug, Lisa M. Komoroske, Brian S. Cheng

## Abstract

Populations across a species’ range may be locally adapted, and failure to recognize this variation can lead to inaccurate predictions of their resilience or vulnerability to climate change. Because life history traits are directly linked to fitness, life history theory can serve as a useful framework for evaluating how populations within species may respond to rapid environmental change. However, relatively few studies quantify multiple life history traits and their tradeoffs across many populations, especially in marine taxa. Here, we used a 10-month laboratory experiment to quantify a suite of reproductive traits in populations spanning the strongest latitudinal temperature gradient in the world’s coastal oceans. We examined reproductive traits in wild-captured adults exposed to simulated local conditions for 7 native Atlantic and 4 introduced Pacific populations of the marine predatory gastropod *Urosalpinx cinerea*. Our data reveals that reproductive season length, the number of reproductive attempts, and annual fecundity unimodally peaked at mid-latitude populations, the species’ range-center. Introduced populations had comparably few spawning attempts and low fecundity despite a longer reproductive period in a less seasonal environment. We then conducted a second experiment quantifying thermal tolerance of developing embryos from 3 native populations, which revealed high sensitivity to temperature at early life stages but weak population differentiation. Taken together, our data reveal stark differences in reproduction that appear to reflect “fast” and “slow” paced lifestyles, which may maximize fitness by spreading the risk of reproductive failure over a single season or lifetime. Our results indicate that warm range-edge populations are highly vulnerable to warming, as low embryonic thermal tolerance may shorten the spawning season and warming is likely to reduce fecundity. This study highlights heterogeneity in life history traits across marine populations that may underlie differential vulnerability to climate warming.

**Open research statement:** All data and code will be publicly available via Figshare and the NSF Biological and Chemical Oceanography Data Management Office (BCO-DMO).

## Introduction

Increasingly, climate change studies have recognized that populations within species are heterogeneous, with physiological performance and life history traits varying substantially across environmental gradients (Villeneuve et al., 2021a, b; Sasaki et al., 2022). Consequently, forecasting models that ignore trait variation can grossly overestimate population resilience under rapid environmental change (Cacciapaglia & van Woesik 2018; Donham et al., 2023). Since life history traits are phenotypes that directly affect fitness, there is a need to identify environmental drivers of intraspecific trait variation (e.g., Ricklefs & Wikelski 2002). Here, “space-for-time” substitution, which assumes that phenotype-environment matching over space approximates phenotypic change over time, may yield insight into the effects of climate change on life history traits and population persistence into the future (e.g., Lovell et al., 2023). This is particularly valuable for longer-lived taxa where multi-generational laboratory studies of adaptation are impractical or constrained to few populations.

Life history theory predicts that finite energy budgets drive trade-offs in traits like fecundity, longevity, and parental investment in offspring (*fecundity-longevity*, Kirkland & Rose, 1991; Travers et al., 2015; *offspring size-number*, Smith & Fretwell, 1974; Marshall et al., 2008; *offspring number-provisioning*, Clarke & Gore 1992). In any given environment, selection is expected to favor life history strategies that maximize lifetime reproductive success. However, conflicting evidence of how life history traits vary across steep environmental gradients (i.e., with latitude) exists both within and among taxonomic groups (See Appendix: Table S1 for non-exhaustive summary). Moreover, most studies of biogeographical variation in life history focus on species contrasts (e.g., Young, 1990, Marshall et al., 2018). This foundational work has set the stage of life history theory but relatively few studies evaluate within species life history variation, especially for marine taxa. An understanding of how life history traits vary across marine populations is needed because climate velocities are greater in the ocean (Burrows et al., 2011) and because population differentiation can be stronger in marine versus terrestrial systems (Sasaki et al., 2022). Considerable knowledge gaps also remain for entire marine taxonomic groups and developmental modes, with most studies focused on bony fish and crustaceans with highly dispersive larval stages. Despite the greater potential for local adaptation in species with limited dispersal (Sasaki et al., 2022), few studies evaluate life history variation in direct-developing taxa (e.g., Waite et al., 2024).

Theory suggests that at high latitudes, shorter growing seasons and harsh winter conditions are expected to restrict spawning season length, while selecting for longer-lived individuals that have fewer, larger clutches annually (e.g., Conover, 1992; Slesinger et al., 2021). Conversely, at low latitudes, near-constant access to resources and longer growing seasons should relax selection on clutch size and number across comparatively longer spawning periods, selecting for multiple smaller clutches annually. This continuum of fast to slow pace-of-life (POL) describes the evolution of life history strategies prioritizing adult reproduction (fast-paced) or survival (slow-paced; Promislow & Harvey, 1990). With latitude, POL should reflect patterns in mortality (e.g., predation; Harper & Peck, 2016), the cost of reproduction (Thorson, 1950), and parental investment (Thatje & Hall, 2016). Taken together, classic life history theory predicts trade-offs in offspring number and size (“many, small” or “few, large”; e.g., Smith & Fretwell, 1974), and between offspring number and provisioning (e.g., Clarke & Gore, 1992). Further complicating matters, reproductive traits that are likely to vary at the clutch level and over a season, like offspring size and provisioning, need not be correlated with annual fecundity, and instead may reflect seasonal constraints on (and variation in) reproductive effort (Smith & Fretwell, 1974). Offspring size and provisioning may also increase with latitude given expected size-specific winter mortality during the first year of life (Pettersen et al., 2020). Relative to native populations, introduced Pacific populations experiencing less seasonal environments are expected to have broader spawning seasons, while retaining similar life history traits.

To address these knowledge gaps, we focus on a widely distributed, direct-developing gastropod, the Atlantic oyster drill (*Urosalpinx cinerea;* hereafter *Urosalpinx*). The native range of this species spans the strongest latitudinal temperature gradient in the world’s coastal oceans, the Atlantic coastline of North America (Baumann & Doherty, 2013), varying in 9°C mean summer temperature and by 2-fold in season length (Villeneuve et al., 2021b). Furthermore, evidence reveals that there is local adaptation in thermal performance among native Atlantic *Urosalpinx* populations (Villeneuve et al., 2021a & 2021b), suggesting the potential for latitudinal differences in reproductive phenology and life history strategies. Genomic analyses have also revealed strong population structure throughout this species’ native Atlantic and introduced Pacific range, indicating limited population connectivity (Bentley et al., 2024). *Urosalpinx* life history is particularly amenable to testing life history theory in the laboratory, as reproductive phenology and life history traits are easily measured at the individual level. *Urosalpinx* females lay transparent, benthic egg capsules, from which trade-offs in offspring number (fecundity), size, and provisioning can be readily quantified.

## Methods

### Experiment 1: reproductive life history traits

To quantify reproductive life history variation in native Atlantic and introduced Pacific populations, we exposed wild collected oyster drills to simulated seasonal changes in seawater temperature using laboratory microcosms. This “observational experimental” approach has the advantage of facilitating fine scale temporal monitoring that would be impossible in a field study, and has the benefit of holding other environmental covariates constant (e.g., food availability, salinity). To balance replication across populations with logistical constraints, we designed a laboratory experiment that grouped populations into 4 regions: 1) “North-Atlantic” native populations were sourced from Rhode Island (RI), Massachusetts (MA), and New Hampshire (NH), 2) “Mid-Atlantic” native populations from Delaware (DE) and North Carolina (NC), 3) “South-Atlantic” native populations from South Carolina (SC) and Georgia (GA), and 4) “Pacific” introduced populations from San Francisco Bay (Coyote Point, CP; Richardson Bay, RB) and nearby Tomales Bay (TO) in California, and from Willapa Bay (WP) in Washington. We then exposed oyster drills within each region to experimental temperature regimes based on a 3-year daily average of seawater temperatures from the nearest subsurface NOAA buoy (NOAA National Data Buoy Center, CO-OPS API) that averaged across all sites within a region. Further details are provided in the supplemental materials.

We collected adult oyster drills from February to June 2023 (See Appendix: Table S2). We based the timing of Atlantic collections on best available phenological records, aiming to precede the onset of spawning (Carriker 1995; *personal observation*). In the absence historical records, we collected adult snails from California and Washington in early June when seawater temperatures (i.e., 12°C) were comparable to the onset of spawning in Atlantic populations. Within 1.5 weeks of collection, we sexed and haphazardly assigned individuals within the size range of 15-30mm shell height to establish breeding pairs for each population. We maintained 20 breeding pairs for all populations except for Delaware (n=12).

We used four recirculating seawater systems (1 for each region) to simulate daily changes in seawater temperature for ten months. Each recirculating seawater system possessed temperature control through inline chillers and titanium heaters, along with foam fractionators, and biological and mechanical filtration. We monitored water quality continuously for temperature (HOBO UA-001-64, Onset, Bourne, MA, USA), daily for salinity (YSI Pro, YSI Incorporated, OH, USA), and twice-weekly for ammonia (RedSea USA, Houston, TX, USA). We performed routine water changes as needed to maintain salinity from 29-31 and ammonia levels below 0.2 ppm. Seawater was produced using Instant Ocean Sea salt and reverse osmosis deionized water. We housed replicate breeding pairs of snails within each seawater system in cylindrical plastic containers (5 × 10 cm, 250 ml volume) retrofitted with two vinyl screen windows (4×8 cm; 1 mm^2^ pore size) to allow for passive flow. Breeding pairs were fed *ad libitum* with 1-2 cm juvenile oysters (*Crassostrea virginica*) replaced weekly and sourced from an aquaculture facility (Atlantic Aquafarms, Gloucester, VA, USA).

To characterize reproductive phenology, we observed breeding pairs every weekday and recorded the onset and cessation of egg laying. We defined a “clutch” as a group of eggs laid by an individual female over consecutive days, with the cessation of spawning occurring when the female physically moved away from eggs or resumed feeding. We measured reproductive output as the total number of clutches, egg capsules, embryos, and juvenile hatchlings produced per female during the experimental period. For each clutch, we counted the number of egg capsules and embryos within each capsule using a stereomicroscope (Leica S9D, Leica Microsystems, Wetzlar, Germany). We then quantified the number of viable hatchlings produced per female by rearing clutches in numbered plastic mesh tea strainers (500-µm mesh pore size, Upton Tea Imports, Holliston, MA) at treatment conditions for the duration of encapsulated development. At the first juvenile emergence, we enumerated and preserved all individuals for later measurement of juvenile hatchling shell size.

We quantified the tradeoff between the number of offspring produced and maternal investment per offspring as the per capita capsule albumen available to developing embryos, or the capsule area mm^2^ embryo^-1^ (Rivest 1983). To measure capsule volume, we photographed all egg capsules within a clutch at a known magnification and standardized orientation. After excluding the capsule stalk in image preprocessing, we measured the capsule area via automated particle analysis in Fiji ImageJ (Schindelin et al., 2012).

We further investigated the tradeoff between the number of offspring and offspring fitness as population differences in juvenile size at hatching and hatching success. We measured juvenile size in recently emerged juveniles as the particle area of individuals photographed aperture down. Shell area was used to minimize potential confounding effects of shell allometry observed in older juvenile and adult *Urosalpinx* (*personal observation*). We finally measured survival to hatching, as the proportion of embryos in a clutch that survived to the hatchling stage.

### Experiment 2: embryonic thermal tolerance

We conducted a subsequent experiment in May to September 2024 to evaluate whether population-level differences in thermal tolerance during early life stages may explain temporal breaks in adult reproduction observed in Experiment 1. Here, we quantified temperature dependent hatching success of F2 laboratory reared individuals sourced from a parallel long-term experiment on *Urosalpinx* (Dwane et al., *in preparation*). Briefly, in May 2022, F0 adults were collected from the same GA, NC, and NH populations as in the present study, but were spawned under common conditions (18°C). In August 2022, F1 offspring from each population were placed in simulated thermal environments from their original collection site and allowed to feed *ad-libitum* for two reproductive seasons. We then evaluated hatching success of F2 embryos produced by the above F1 parents in their second reproductive season. We used a split-clutch design, where egg capsules from each clutch were allocated to seven constant temperature treatments (18, 20, 24, 28, 30, 32, 34°C). The split-clutch design allows us to mitigate confounding effects of female identity (e.g., female size, offspring provisioning) and the timing of reproduction (e.g., temperature at time of laying). Developmental failure was determined by an arrestment in development. Temperature treatments reflect conditions that adults commonly experience (18-30°C) and chronic temperature (32, 34°C) conditions for lower latitude Atlantic populations (Carriker 1955).

### Data analysis

We performed all analyses in R (version 4.4.2, R Core Team 2024), using generalized linear mixed models (R packages ‘*glmmTMB’*; Brooks et al. 2017). For detailed descriptions of statistical models, see Appendix: Table S3. Model assumptions were assessed using *DHARMa* (Hartig, 2023) and *performance* packages (Lüdecke et al., 2020), specifically evaluating residual normality, homoscedasticity, zero-inflation, dispersion, and outlier/influential data points. Fitted GLMMs were assessed using the car package for type II analyses of deviance. When applicable, Tukey post-hoc tests were conducted using the “pairs” and “ghlt” functions from the *emmeans* and *multicomp* packages (Lenth 2024; Hothorn et al., 2008). All models were visualized as estimated population means with 95% confidence intervals. Outliers with high leverage were removed from analyses if standardized residuals were >3.

Due to difficulties with collecting smaller size classes of *Urosalpinx* in Atlantic range-edge populations and a lack of large adults in NC, average adult size significantly differed among populations for both females (GLMM; χ^2^ = 251.18, df = 10, p < 0.001) and males (χ^2^ = 358.85, df = 10, p < 0.001). Female snails were on average larger in GA, SC, and NH, and as well as TO and WP in the Pacific range (See Appendix: Figure S3 for adult size distributions). To explore the potential effect of female size on response metrics, all statistical models were constructed with the full and a restricted dataset (15 to 25 mm shell height) and compared for model fit and congruence in population-level effects. Models using size-restricted data still revealed significant differences in average female size among populations (χ^2^ = 33.6, df = 7, p < 0.001) and resulted in three populations becoming data deficient (NH, TO, WP). Nevertheless, models using the size-restricted dataset produced similar latitudinal patterns to those with the full dataset. Therefore, all populations and biological replicates were retained in subsequent analyses, with caveats related to female size noted when applicable.

## Results

### Reproductive phenology

In Atlantic populations, we identified three distinct patterns of reproductive phenology that varied latitudinally (Fig. 1). At mid-latitudes, the species range center, we observed near continual spawning in all NC females over an 8-month period and upwards of 13 clutches laid per individual. In contrast, repeated laying at low latitudes was constrained to two pulses, where a multi-month hiatus in spawning coincided with the warmest annual sea water temperatures and at most 7 clutches laid per individual. Although similar temporal gaps in the spawning were observed in most high latitude populations, NH females exclusively spawned once either early or late in the season. We note that while most high- and mid-latitude Atlantic populations had comparably sized individuals, NH females were on average larger (Appendix: Fig. S3). Among Atlantic populations, MA (15%, n=20) and NH (40%) had the lowest proportion of reproducing females, while nearly all RI (85%) and NC (100%) spawned at least once during the experimental period. Surprisingly, the high-latitude RI population displayed a single pulse pattern early to mid-season with highly variable numbers of clutches per female (Appendix: Fig. S4d). Delaware individuals, that experienced warmer (+3°C ambient) conditions in this experiment than in the field, had variable phenological patterns and number of clutches irrespective of female size, and including singleton, continuous, and early/late season spawning.

**Figure 1.**
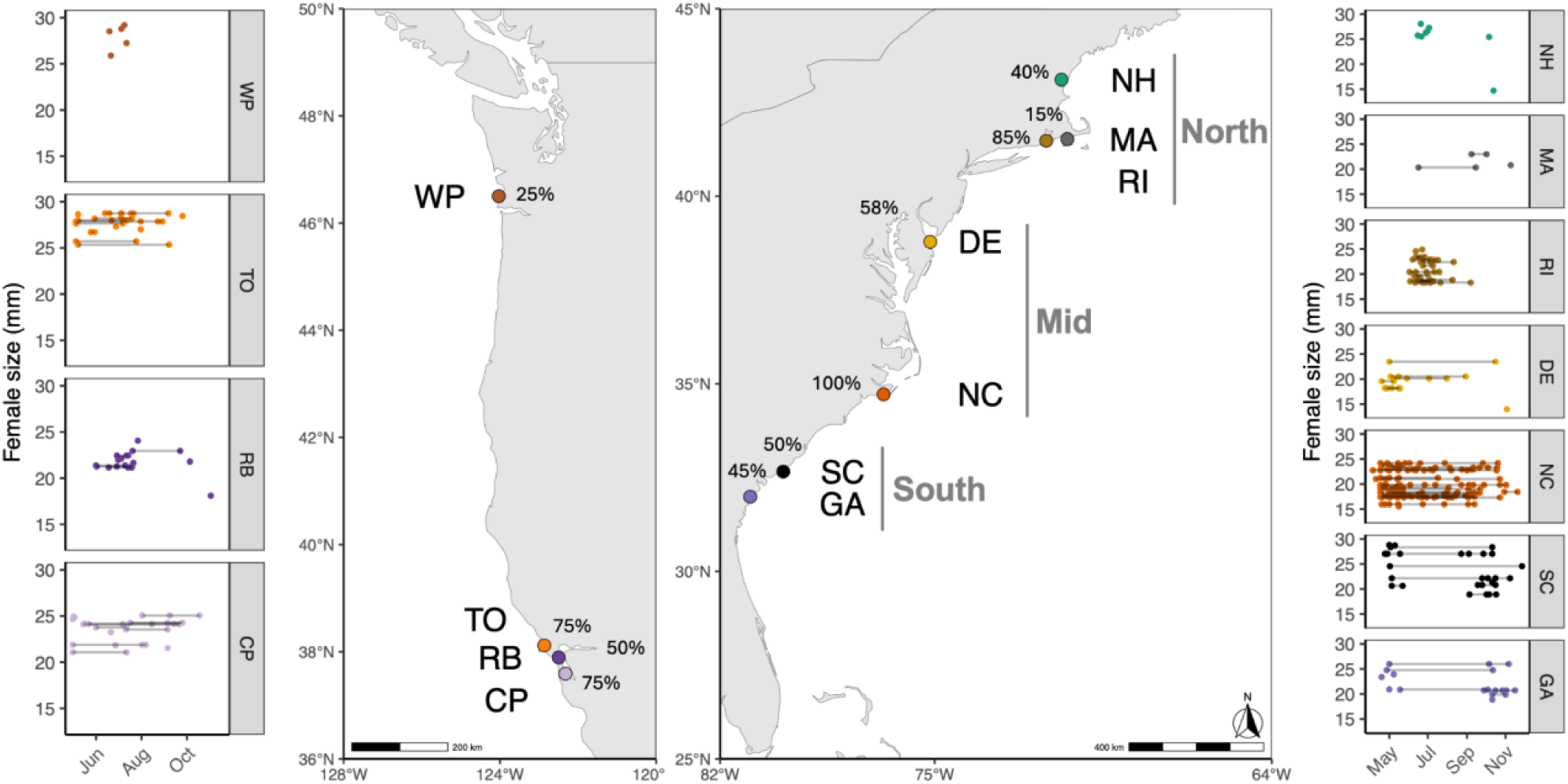
Collection sites and phenology and frequency of reproductive output from Experiment 1. Scatterplots illustrate size-specific reproductive phenology of introduced Pacific (left) and native Atlantic (right) *Urosalpinx* in the coastal USA. To balance sample size and experimental constraints, reproductive adults were experimentally grouped as indicated by vertical lines on the Atlantic coast (all Pacific populations were grouped together). Within plots, gray horizontal bars represent the start and end of the spawning period for an individual female, while points denote discrete clutches. Points without bars represent females that laid only once. Inset percentages represent the proportion of females from each population that spawned at least once. Vector maps of US coastlines were sourced from the “rnaturalearthdata.” Note the difference in spatial scales for the two maps.

As in the native species range, reproductive phenology differed regionally in introduced Pacific populations. We observed similar patterns between high-latitude Pacific (WP) and Atlantic (NH) populations, with females laying singleton clutches, but exclusively early in the season (Fig. 1). In contrast, low-latitude California populations displayed intra-population variation in the timing of spawning, (Fig. 1) but not the average number of clutches produced per female (See Appendix: Fig. S4a; Tukey-Kramer, p < 0.05).

### Fecundity

There was up to an order of magnitude difference in reproductive output across Atlantic populations (Fig. 2). Furthermore, we observed that reproductive output unimodally peaked at the native range center and declined towards the range edges. This was consistent across all metrics: total number of embryos (χ^2^ = 100.3, df = 10, p < 0.001; Fig. 2d), clutches (χ^2^ = 138.5, df = 10, p < 0.001; Appendix: Fig. S4d), egg capsules (χ^2^ = 129.7, df = 10, p < 0.001; Fig. S4e), and juvenile hatchlings per female (χ^2^ = 120.9, df = 10, p < 0.001; Fig. S4f). High-latitude NH and MA consistently had the lowest reproductive output relative to all other Atlantic populations (See Fig. 2d and Appendix: Fig. S4b,e,f; Tukey-Kramer, p<0.05). Reproductive output was comparably low in the high-latitude Pacific (WP) relative to Atlantic populations (Fig. 2a,d; Appendix: Fig. S4). A similar pattern was also observed among Pacific populations, with reproductive output being significantly lower in high-latitude WP relative to all low-latitude California (CP, RB, TO) populations (Appendix: Fig. S4a,b,c; Tukey-Kramer, p < 0.05). Despite differences in phenology, Pacific populations had similar reproductive output to range-edge Atlantic populations (See Fig 2a and Fig. S4a,b,c; Tukey-Kramer, p > 0.05).

**Figure 2.**
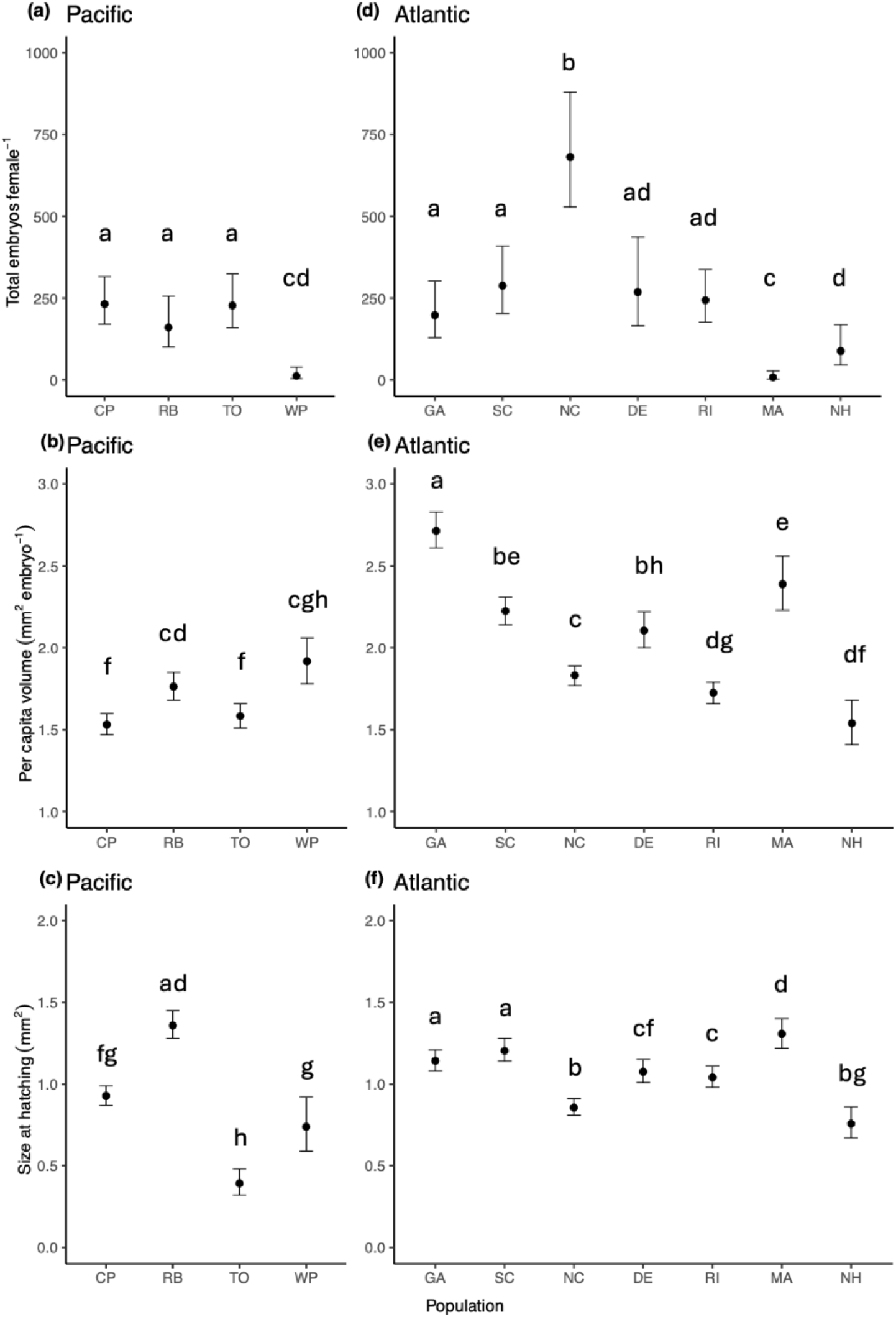
Average number of embryos laid per female (a,d), per capita capsule volume (b,e) and offspring size at hatching (c,f) ± 95^th^ percentile confidence intervals in Atlantic and Pacific populations (south to north from left to right on the x-axis). Letters denote significant pairwise differences.

The reproductive output of embryos also varied significantly with female body size, and that this relationship was strongest in Atlantic mid-latitudes (Female size, χ^2^ = 7.35, df = 1, p < 0.001; Population x Female size, p = 0.33, Fig S5a). Interestingly, female size was a significant predictor of reproductive output of hatchlings (Female size, χ^2^ = 11.4, df = 1, p < 0.01; Population x Female size, χ^2^ = 21.4, df = 10, p = 0.02), but not the number of clutches (Female size, χ^2^ = 0.68, df = 1, p =0.4; Population x Female size, p = 0.96) or capsules laid per female (Female size, χ^2^ = 0.72, df = 1, p =0.4; Population x Female size, p = 0.68).

### Offspring provisioning and fitness

Despite a mid-latitude unimodal peak in reproductive output, offspring provisioning declined with increasing latitude in Atlantic populations (Population, χ^2^ = 2290.1, df = 10, p < 0.001; Fig. 2e) irrespective of female size (χ^2^ = 3.7, df = 1, p = 0.054; Population x Female size, χ^2^ = 517.6, df = 10, p < 0.001). Relative to Atlantic populations, Pacific populations had comparatively low offspring provisioning, with the least provisioning in TO and CP (Fig 2b). Population differences in offspring provisioning were, in part, explained by a latitudinal decline in egg capsule size from GA to RI, but not in embryo density (See Appendix: Fig. S6d for embryo densities and Fig. S6e for capsule size). Embryo densities were similar among most Atlantic populations (excluding NH, ∼6-8 embryos capsule^-1^) and on average higher in Pacific populations (Fig. S6a,d; Population, χ^2^ = 1334.3, df = 10, p < 0.001; Female size, χ^2^ = 279.0, df = 1, p < 0.001; Population x Female size, χ^2^ = 133.3.9, df = 10, p < 0.001). We note that deviations existed in high-latitude populations: low provisioning in NH was driven by anomalously high embryos densities while high provisioning in MA was driven by intermediate sized capsules with low embryo densities. Although capsule size varied significantly among populations, female size had a significant positive direct and interactive effect on capsule size (Population, χ^2^ = 1038.2, df = 10, p < 0.001; Female size, χ^2^ = 1416.8, df = 1, p < 0.001; Population x Female size, χ^2^ = 190.5, df = 10, p < 0.001).

Offspring provisioning was positively correlated with offspring size at hatching, but the strength of this relationship varied greatly by population (Fig. S5b). Although there was no significant association between offspring size and provisioning in NC (p = 0.19), hatchlings were ∼20 to 30% smaller in size in NC relative to SC, MA, and RI. In contrast, we found that GA hatchlings had ∼ 25% greater provisioning than in SC, but were similar in size. Overall, hatchling size varied significantly by population (Fig. 2f; χ^2^ = 1093.4, df = 10, p < 0.001) and female size (χ^2^ = 184.4, df = 1, p < 0.001; Population x Female size, χ^2^ = 273.9, df = 10, p < 0.001), with juveniles being on average smaller in NC relative to all other Atlantic populations.

In the first experiment, survival of F1 embryo to the hatchling juveniles lifestage significantly differed among populations (Appendix S1: Fig. S6c, S6f; Population, χ^2^ = 37.5, df = 10, p < 0.001; Female size, χ^2^ = 1.4, df = 1, p =0.23; Population x Female size, χ^2^ = 16.4, df = 10, p = 0.09). Although few significant pairwise differences existed among Atlantic populations, all populations had greater than 60% survivorship. We observed that survivorship was on average highest in MA, but significantly lower in compared to the other high latitude populations (i.e., NH and RI, Tukey;Kramer; p < 0.05). Similarly, survival to hatching was lowest in WP (∼30%) relative to all other Pacific populations (55-75%).

### Experiment 2: embryonic thermal tolerance

Survival of F2 offspring varied significantly among GA, NC, and NH populations (Fig. 3; χ^2^ = 10.5, df = 2, p = 0.005), and decreased with increasing temperature during encapsulated development (χ^2^ = 187.1, df = 1, p < 0.001). Significant pairwise differences in hatching success were observed between NH and NC populations, but only at temperatures exceeding 24℃ (Tukey-Kramer: p_24℃_ = 0.03, p_28℃_ < 0.001, p_30℃_ < 0.001, p_32℃_ = 0.002, p_34℃_ = 0.003).

**Figure 3.**
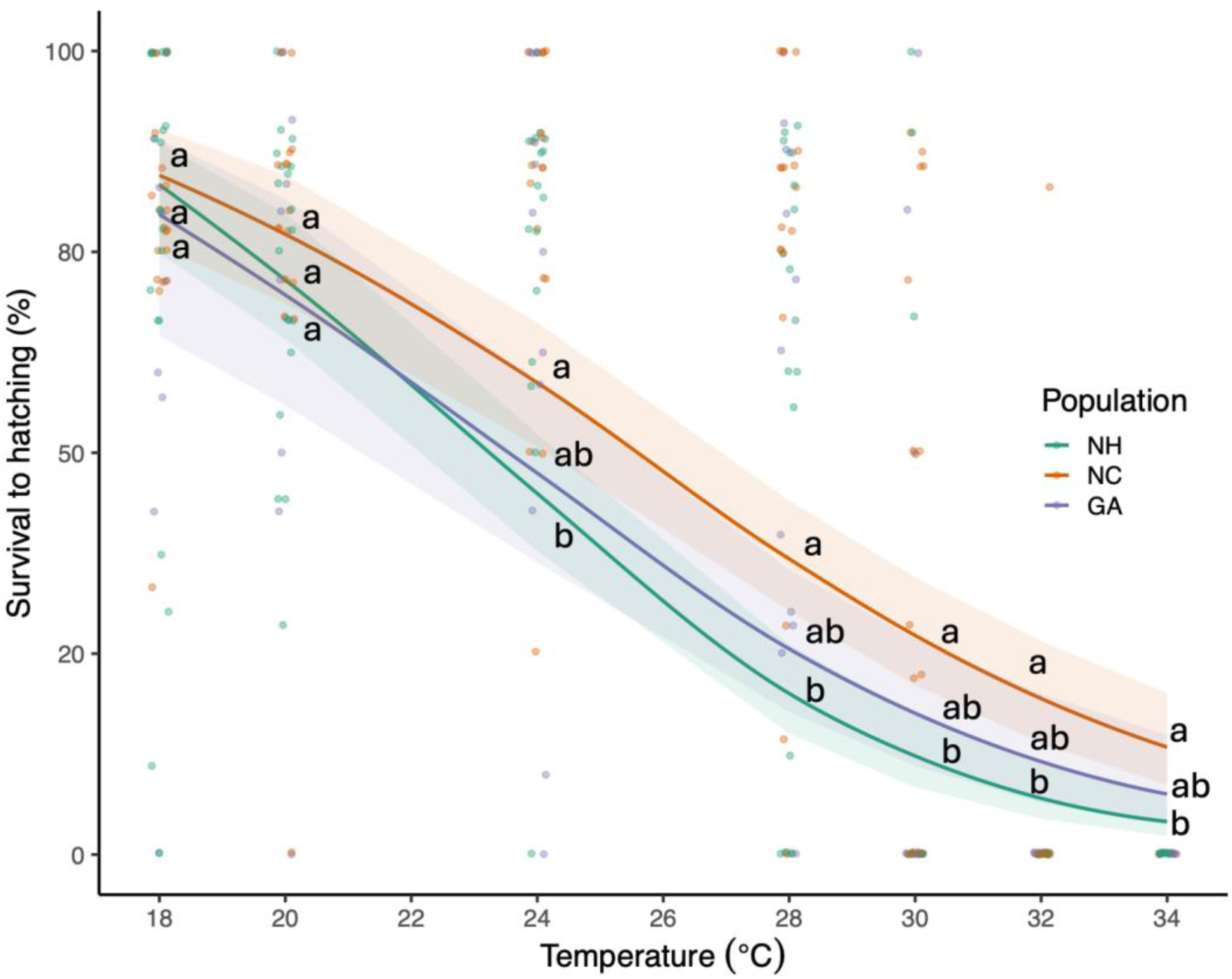
Embryonic thermal tolerance across populations from Experiment 2. Each point represents survival to hatching, defined as the percentage of embryos from a single egg case that hatched (± 95th percentile confidence intervals) in F2 laboratory reared offspring under constant temperature conditions. Letters denote significant pairwise differences among populations at discrete experimental temperatures: 18, 20, 24, 28, 30, 32, and 34°C (Tukey-Kramer, p<0.05). Points are slightly jittered for clarity.

## Discussion

We observed strong divergence in reproductive phenology, fecundity, and offspring performance across the native and invasive range of *Urosalpinx* spanning 11° of latitude. These population-level differences in reproduction likely reflect alternative life history strategies that maximize fitness by spreading the risk of reproductive failure over a single reproductive season (e.g., at mid-latitudes) or a lifetime (at high- and low-latitudes). While we observed the classic tradeoff of mothers investing in “many, small” or “few, large” offspring, we surprisingly found steep declines in offspring provisioning with increasing latitude that were not associated with declines in offspring size. Overall, our findings conflict with theoretical expectations of linear monotonic variation in life history traits across latitude, revealing non-linearities in reproduction, possibly arising from environmental seasonality combined with additional drivers of adult and juvenile mortality rates (e.g., predation, food availability).

### Reproductive phenology

Contrary to expectations of reproductive season length and clutch number decreasing with increasing latitude and seasonality (e.g., Conover 1992), we observed unimodal peaks in both traits at mid-latitudes. Latitudinal patterns in these traits were not monotonic, and therefore were inconsistent with empirical studies of inter- and intra-specific life history variation (See Table S1). However, observations of the shortest reproductive seasons and fewest spawning attempts in high latitude populations do align with theoretical expectations. Fewer and on average larger females reproduced at the highest latitude population, suggesting that cooler, seasonal environments with shorter growing seasons may select for reduced annual investment in reproduction and/or delayed reproductive maturity (Huret et al., 2019; Álvarez-Noriega et al., 2023). At high latitudes, shorter growing seasons likely confer smaller and/or more restricted metabolic energy budgets, and therefore reproductive failure may incur a higher fitness cost relative to populations with larger, more flexible energy budgets at lower latitudes (Schultz & Conover 1997; Huret et al., 2019). Therefore, high-latitude populations should experience increased selection against reproduction when the likelihood of failure is predictably high (Jourdan-Pineau et al., 2022). Emergent patterns in spawning phenology may reflect parent-offspring conflict over the optimal timing of spawning, such that adult females and yearling juveniles have adequate time for growth and energy storage in advance of winter (Middaugh & Hemmer, 1992; Varpe, 2017).

In contrast to the classical model of long growing seasons supporting continuous reproduction (e.g., van de Kerk et al., 2016), we observed two-pulse reproduction in low latitude environments. This aligns with recent simulation-based models that found reproductive seasonality as favorable in highly predictable environments, as well as those with high environmental seasonality and low productivity (Burtschell et al., 2023). Here, physiological constraints on early and adult life stages during the warmest period of the year may explain a two-pulse, or early- and late-season spawning pattern. Temperature thresholds for embryological developmental failure are associated with a cessation in adult spawning (LT_50,GA_ = 24°C). This strong seasonal variation in offspring value, or the potential contribution to fitness, may explain why mothers skip spawning at times when resources do not appear to be limiting (Varpe, 2017). As in bivalve molluscs across environmental latitudinal gradients, the two-pulse reproductive patterns in *Urosalpinx* may synchronize offspring recruitment with peak food availability and optimal thermal conditions (Philippart et al., 2014; Oyarzún et al., 2018).

Overall, divergence in reproductive phenology of mid- and range-edge populations best reflect “fast” and “slow” POL, which differentially balance the cost of current reproduction with long-term survival (Stearns, 1992; Moss et al., 2017). These lifespan-fecundity tradeoffs are, in part, corroborated by a parallel long-term experiment on laboratory reared F1 *Urosalpinx* which revealed that reproduction during the first year of life is common in NC and rare in NH populations (Dwane et al., *in preparation*). Higher fecundity at smaller body sizes in mid-latitude females suggests a “fast” POL, prioritizing reproduction earlier in life to maximize fitness across a comparatively short lifespan (Stearns, 1992). Higher shell-crushing predation rates at mid-latitudes may select for faster POL (Harper & Peck, 2016). Conversely, in high-latitude populations, slow-paced but longer-lived individuals may maximize lifetime fitness by skipping breeding opportunities (Audzijonyte & Richards, 2018) and reproducing at larger body sizes to realize higher fecundity (e.g., Stearns & Crandall, 1981; Álvarez-Noriega et al., 2023).

### Fecundity

We observed unimodal peaks at mid-latitudes for clutch size and total reproductive output per female. Despite expectations of the cost of reproduction declining with increasing season length and temperature at lower latitudes (Sletvold & Ågren, 2015), low fecundity occurred at both low- and high-latitudes. Unimodal patterns in fecundity suggest that performance declines towards a species range edge (e.g., ‘abundant center hypothesis’ *sensu* Sagarin & Gaines, 2002), although local selective pressures could also create this pattern (e.g., food abundance, Lester et al., 2007). We posit that annual differences in the number of clutches and fecundity between mid-Atlantic populations and all other Atlantic and Pacific populations represent income and capital breeding strategies, respectively (e.g., Jönsson, 1997; Sainmont et al., 2014). Income breeding strategies are only viable in environments with continual access to food, as continual reproduction is financed by rapid turnover of assimilated energy (i.e., the “income”) *in lieu* of energy storage (Varpe et al., 2009). In contrast, capital breeding is favorable in seasonal environments where food is predictably limiting over time and stored energy, or “capital,” is necessary to balance routine maintenance costs (e.g., overwintering) with present and future reproduction. Like “fast-slow” POL, populations employing income and capital breeding strategies should exhibit tradeoffs in longevity and lifetime fitness (Wilbur & Rudolf, 2006). Under this framework, we may expect mid-latitude individuals that disinvest in energy storage to be both shorter-lived and more susceptible to environmental stress.

### Life history trade offs

While we observed a weak association between fecundity and offspring size, tradeoffs in fecundity and provisioning were not consistent across *Urosalpinx* populations. This suggests that fecundity alone may be an incomplete proxy for fitness. Instead, optimal offspring size and provisioning strategies likely vary by environment (Smith & Fretwell 1974), with offspring size-performance relationships influenced by temperature (e.g., Fischer et al., 2003), food availability (e.g., Bashey, 2006), intraspecific competition (e.g., Allen et al., 2008), and predation pressure (e.g., Allen, 2008). In combination with life history traits at the adult stage (i.e., reproductive phenology and fecundity), latitudinal patterns in offspring life history traits may provide insight into dominant ontogenetic and spatial drivers of fitness.

Although offspring size is expected to increase with latitude in marine taxa (e.g., Marshall et al., 2018), offspring size was smallest at both mid-latitudes and in the highest latitude population. Investing minimally in many offspring has been associated with unpredictable or variable environments, where spreading the risk of reproductive failure across many individuals is favorable (i.e., high intrinsic offspring mortality, Pianka 1970). Conversely, strategies to produce “few, larger” offspring in low-and most high-latitude populations, may result in comparatively robust individuals (e.g., Marshall et al., 2018; Salas-Yanquin et al., 2021), that are better able to withstand abiotic perturbations or food scarcity (e.g., Einum & Fleming 2004).

Here, we demonstrate that despite declines in provision with latitude, “embryo packing” in the high latitude NH population may, in part, compensate for low reproductive effort, or few clutches and egg capsules annually, without measurable effects on early offspring survival. Size-specific winter mortality should select for greater offspring provisioning at high-latitudes, resulting in larger sizes at hatching with lower mass-specific metabolic costs, or fueling rapid growth in advance of adverse conditions (Pettersen et al. 2020). Instead, in NH *Urosalpinx*, high embryo densities with low offspring provisioning may be accommodated via higher conversion efficiency (as observed in fish; Present & Conover, 1992). Furthermore, developing embryos may disproportionately invest in tissue growth over energy storage to attain larger sizes at hatching (Pettersen et al., 2023).

A critique of the approach used here is that field-collected adults are of unknown age and environmental background, such that latitudinal patterns may be confounded by ontogeny (e.g., age-specific growth rates and reproductive output) or latent effects of environmental stress. For *Urosalpinx*, historical field records suggest that phenological patterns at mid- and high-latitudes are consistent over evolutionarily relevant timescales (i.e., continuous spawning in NC, early- and late-season spawning in MA; Carriker 1955). Additionally, phenological patterns observed in the present study are consistent with those of laboratory reared F1 *U. cinerea* that were reared to maturity over 24 months and descended from the same GA, NC, and NH populations (Dwane et al., *in preparation*). While laboratory studies can minimize confounding effects of unpredictable environmental variation and food limitation, they remain approximations of intraspecific life history variation and phenology-fitness relationships in the field. Field investigations are still needed to validate whether laboratory observations are representative of natural populations, particularly in the context of relative predation rates and food availability.

## Conclusions

Warm range-edge populations may be particularly susceptible to climate warming given their phenology and life history strategy of investing in “few, large” offspring. With continued ocean warming, we expect low latitude populations to have reduced recruitment because of higher developmental failure across fewer clutches, and a narrower spawning window. Because habitat temperatures are close to their lethal limits, these populations are most at risk of local extinction. The limited dispersal potential also suggests that *Urosalpinx* may not be able to colonize poleward habitat, resulting in range contraction instead of range shift.

Our ability to forecast climate impacts hinges on accurate estimates of fitness at the population level. Bet-hedging strategies employed in unpredictable or seasonal environments best explain latitudinal patterns in life history traits in the present study (Marshall et al. 2008; Olofsson et al. 2009). Our findings emphasize the importance of framing life history variation as a hierarchy of tradeoffs with different, and often conflicting, selective pressures operating across life stages. But taken together, this study highlights how key reproductive traits can vary substantially among marine populations and are consequential for forecasting population persistence, shifting life history strategies, and resilience to rapid environmental change.

## Acknowledgements

We thank Angie Korabik, Corryn Knapp, Adam Mims, Blair Bentley, Andrew Villeneuve, and Chris Dwane for field collections. For animal care, we thank Ryan Horrigan, Ana Stucker, Chance Yan, Mikayla Newbrey, Elizabeth Clark, Jordanna Barley, Mary Rutter, Chris Dwane and Liam McCarthy. We also thank Chris Smith for oyster seed.

## Funding source

This work was supported by NSF BIO-OCE #2023571.

## Author contributions

AR led study conceptualization, methodology, data curation, analysis, and writing. ER, EB, and AS contributed to data collection, data curation, and writing. LK contributed to conceptualization, writing, and funding acquisition. BC contributed to conceptualization, methodology, analysis, writing, and funding acquisition.

## Conflict of interest

The authors have no relevant financial or non-financial interests to disclose.

## Notes

### Competing Interest Statement

The authors have declared no competing interest.

