## Supplementary Figures for "Fast and slow-paced reproductive life history across native and invasive populations of a predatory marine snail"

**Appendix**

**METHODS**

**Experiment 1:**

All NOAA buoys used for calculating population temperature regimes were located within 10 km of collection sites at 0.7-2.7 m water depth. We collated this publicly available seawater temperature data at a population level using the three most recent years of data, constrained to 2019-2022 (See Appendix S1: Table S1 for NOAA Station IDs; Fig. S1 for site-level temperature regimes). We then computed a regional 3-year daily average of seawater temperature by averaging across populations within a geographic region (See Fig. S2). For practical reasons, Delaware and Washington populations were exposed to Mid-Atlantic and California regimes, respectively, and those site-specific temperatures were not included in regional temperature calculations. Since exposure to a warmer (i.e. Mid-Atlantic, +3°C) rather than cooler (i.e. North-Atlantic, -3°C) thermal regime is more informative with continued ocean warming, we decided to include Delaware in the mid-Atlantic region. While we acknowledge annual thermal minima differ between Washington and average California conditions (Δ3°C), these populations experience similar maximum and average temperatures during the four warmest months of the year (June to October) when we expected spawning to take place.

Females and males were paired such that there was marginal ($p_{WP} = 0.051$) to no correlation in body size by sex ($p_{GA} = 0.63$; $p_{SC} = 0.90$; $p_{NC} = 0.44$; $p_{DE} = 0.38$; $p_{RI} = 0.45$; $p_{MA} = 0.40$; $p_{NH} = 0.40$; $p_{CP} = 0.64$; $p_{RB} = 0.66$; $p_{TO} = 0.72$).

We maintained oysters via daily batch feeding of Shellfish diet 1800™ at target levels of 0.4 mg dry weight algae oyster$^{-1}$ (Reed Mariculture, Campbell, CA; Helm and Bourne 2004). Oysters were maintained in a separate recirculating seawater system at constant temperature (18°C). To minimize disturbance of actively laying females, we avoided any handling of experimental jars, such as replacing consumed oysters or removing eggs, until females had completed spawning.

By using per capita capsule volume as a proxy for maternal provisioning we assume that, in the absence of nurse eggs or utilization of embryo yolk, albumen (perivitelline fluid) is the dominant energy source and supply of critical structural lipid precursors in developing gastropod embryos (Rivest 1983; Wijsman and van-Wijck-Batenburg 1987; Stockmann-Bosbach and Althoff 1989; Rivest 1992; Heras et al. 1998; Brante et al. 2009; Bigatti et al. 2014; Han et al. 2021).

**Statistical models**

All models treated female size and population as fixed effects. Population-level differences in total reproductive output were assessed using zero-inflated mixed effects models with negative binomial distributions and where the occurrence of zeroes varied by population (i.e., ziformula = ~ Population). To reduce effects of heteroscedasticity as a function of population, dispersion parameters (i.e. dispformula = ~ Population) were implemented in models of offspring provisioning, offspring size, offspring survival, capsule size, and the number of embryos per capsule (hereafter mentioned as embryo density). For the aforementioned models, mother and clutch were included as random hierarchical effects to account for repeated reproduction by mothers over time and non-independence of repeated measurements within a given clutch (e.g. offspring size). Differences in the offspring provisioning and offspring size were modeled as log normal distributions, while capsule size and embryo density were modeled with gamma and generalized Poisson distributions, respectively. Hatching success of F1 offspring from field collected adults were assessed using zero-inflated mixed effects models with a beta-binomial distribution weighted by the total number of embryos in that clutch. Similarly, hatching success of F2 offspring from laboratory reared adults were modeled with beta-binomial distributions weighted by total number of embryos in that clutch, but allowing the occurrence of zeroes to vary by population. Acknowledging that differences in parental temperature at the time laying may influence F2 egg quality and/or offspring thermal tolerance, we specified parental temperature as a covariate and mother as a random effect.

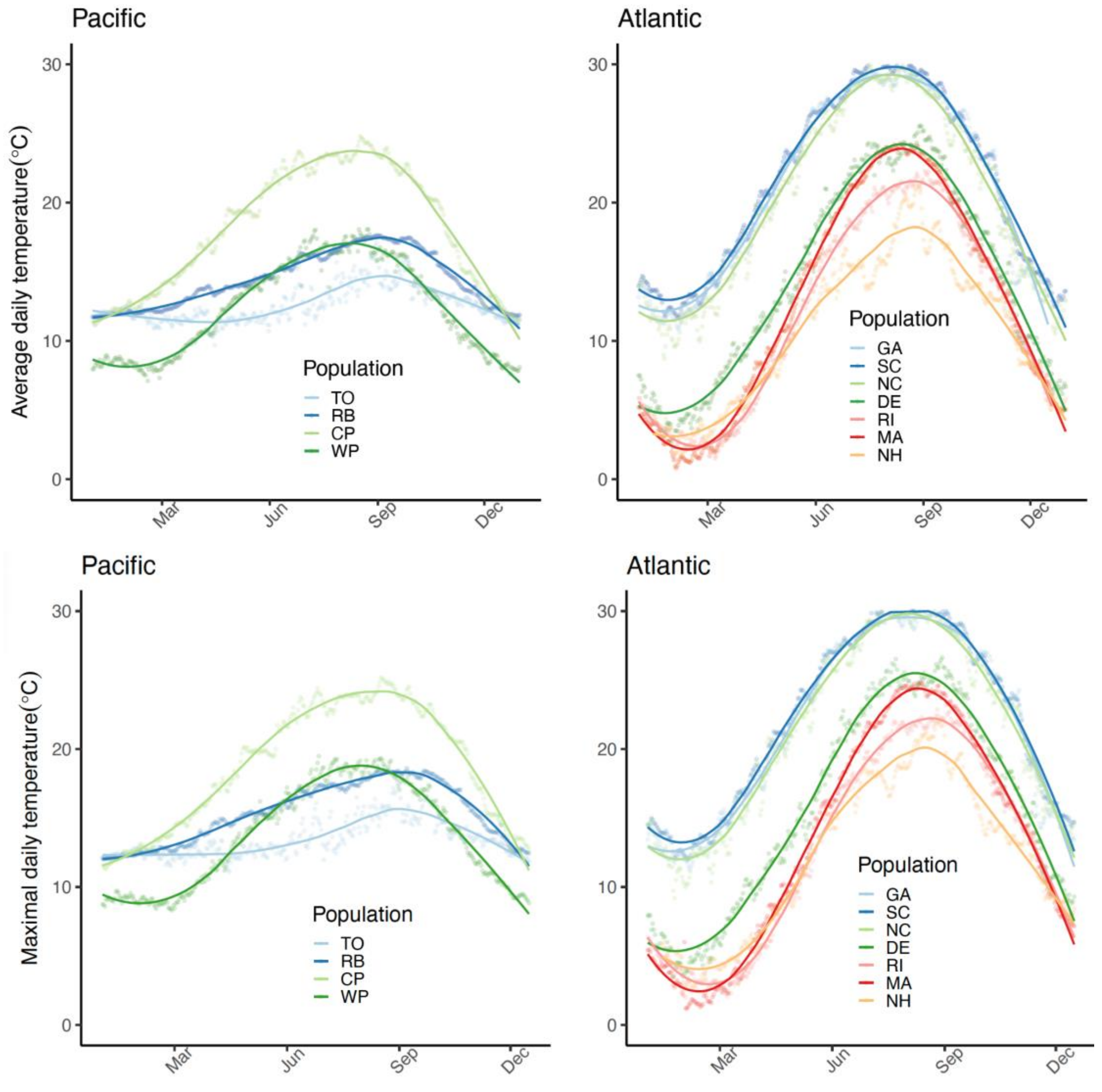


Figure S1. Average daily temperature as determined from most recent 3-year daily averages (loess curve; 2019-2022), where overlayed points are calculated daily average temperatures.

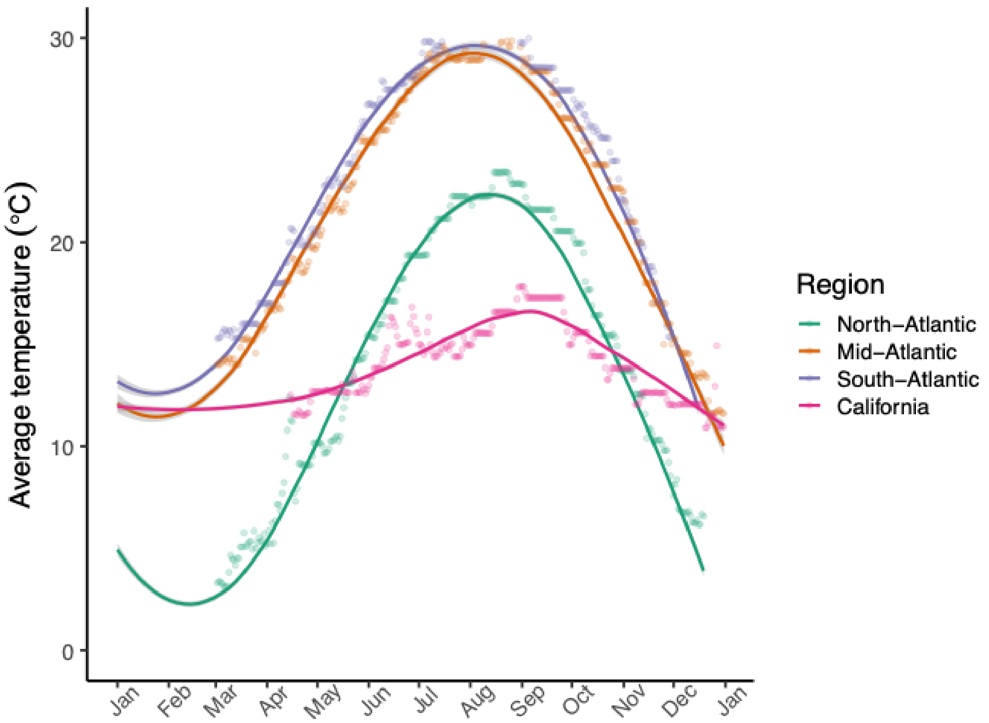


Figure S2. Regional sea surface temperature determined from 3-year daily averages (loess curve; Data source: NOAA CO-OPS API, 2019-2022; See Supplementary Table 1 for site IDs). Overlayed points are semi-continuous (15-minute interval) experimental temperatures as measured by HOBO temperature pendants.

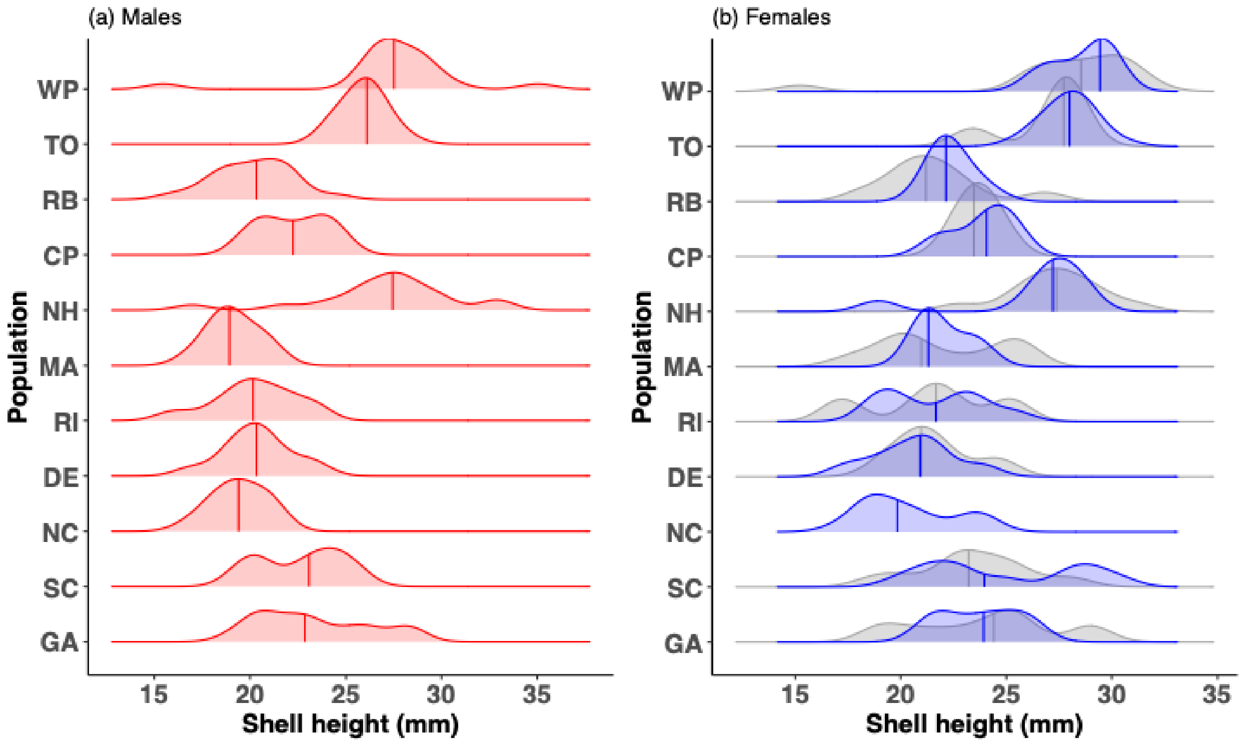


Figure S3. Experimental adult size distributions across populations for (a) males and (b) females, where population medians are represented by vertical bars within ridgelines. Females that spawned at least once or never during the experimental period are represented by grey and blue ridgelines, respectively. NH females were on average larger: $\bar{x}_{NH} = 27.0 \pm 0.6$, $\bar{x}_{MA} = 22.0 \pm 0.6$, $\bar{x}_{RI} = 21.5 \pm 0.5$, $\bar{x}_{DE} = 21.0 \pm 0.6$, $\bar{x}_{NC} = 20.4 \pm 0.5$ mm.

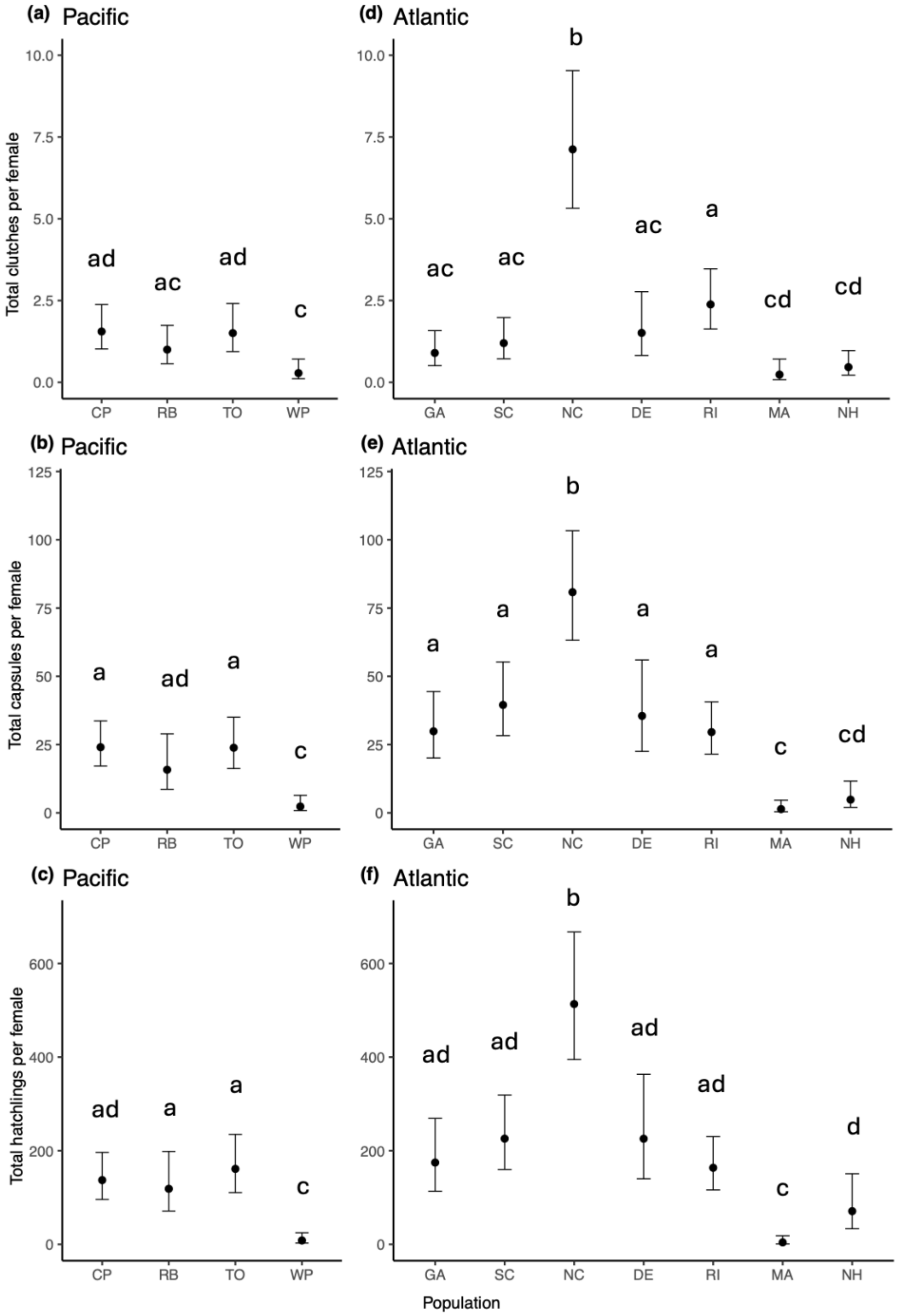


Figure S4. Total reproductive output as represented by (a, d) average number of clutches, (b,e) average number of embryos, and (c,f) average number of hatchlings per female ± 95$^{th}$ percentile confidence intervals in Atlantic and Pacific populations. Letters denote significant pairwise differences (Tukey-Kramer, $p<0.05$).

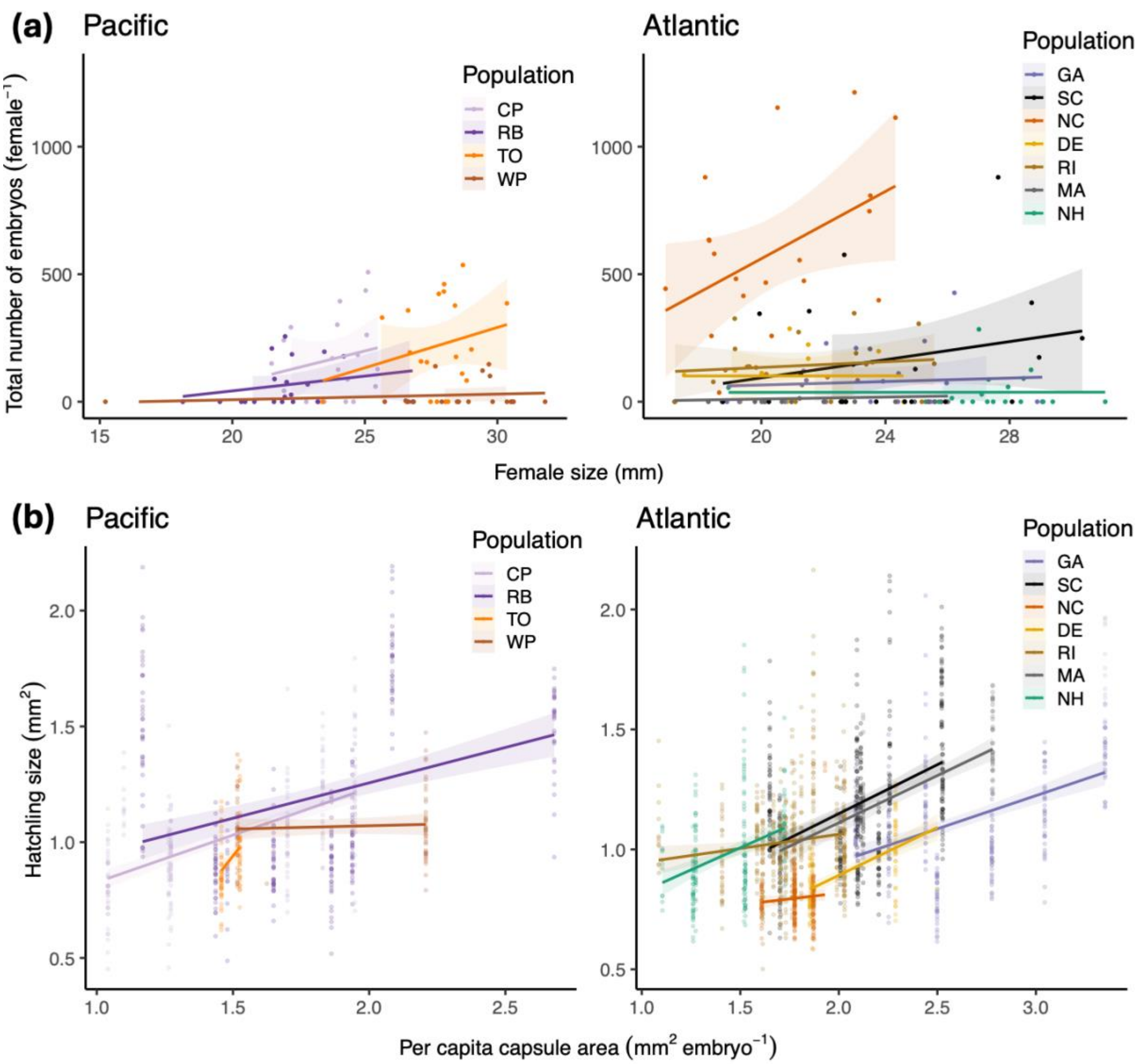


Figure S5. (a) Total number of embryos produced per female as a function of female size (n=12-20). Discrete points are biological replicates. (b) Hatchling size ($mm^2$) relative to average provisioning (per capita capsule area; mm2/embryo) for that clutch. Discrete points represent individual hatchlings, with a subset of 20 hatchlings measured per unique clutch. For clutches with fewer than 20 hatchlings, all individuals were measured. Linear regressions include 95% confidence intervals.

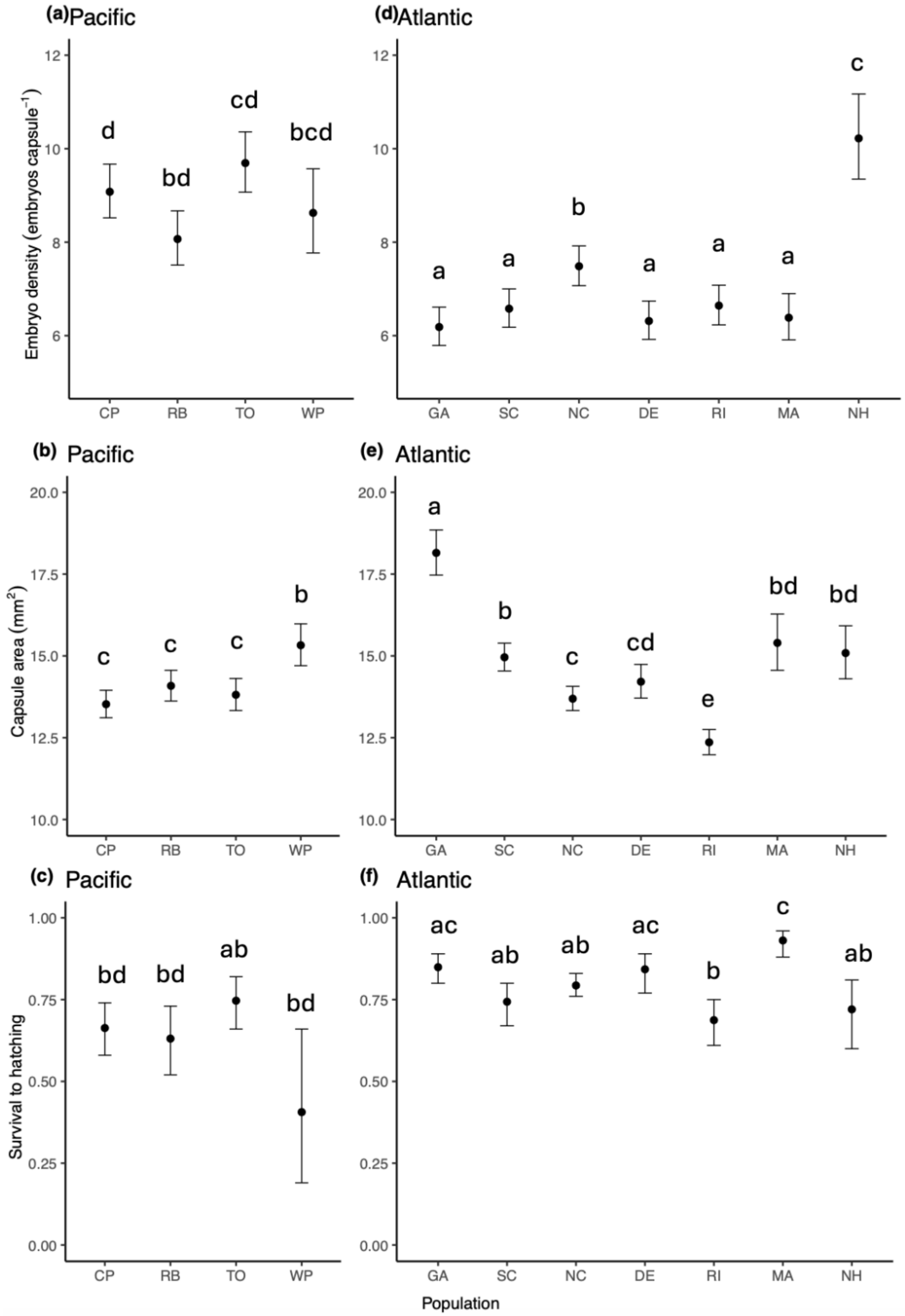


Figure S6. (a,d) Average number of embryos per capsule Average capsule area, (b,e) Capsule area, and (c,f) Embryo survival to hatching ± 95$^{th}$ percentile confidence intervals in Atlantic and Pacific populations. Letters denote significant pairwise differences (Tukey-Kramer, $p<0.05$).

**Table 1.** Theoretical and empirical patterns of reproductive phenology, reproductive output, and offspring performance with latitude (Low = Equatorial or Subtropical; High = Temperate or Polar) or environmental seasonality (low to high) in marine taxonomic orders. Studies on anadromous fish (†) or terrestrial taxa (‡) are noted separately. Estimates of fecundity that were measured at the clutch level in iteroparous species (e.g. gonadosomatic index) are represented as clutch size. Offspring provisioning measurements were proxied by egg volume and diameter. Latitudinal patterns in traits are denoted as linear positive (+), linear negative (–), no correlation (ns), or unimodal (*).

| **Trait** | | **Literature** | **Present Study** | **Interspecific variation** | **Intraspecific variation** | | | |
|---|---|---|---|---|---|---|---|---|
| | | *Low → High* | *Low → High* | | Chordata | Arthropoda | Mollusca | Other |
| **Reproductive phenology** | Season length | 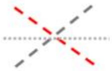 | 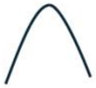 | (–) 301 fish spp.[42]<br>(–) 11 Decapod spp.[43]<br>(–) Sygnathiformes spp.[44]<br>(–) Balanomorpha spp.[45] | (+) Scombriformes[29]<br>(–) Acanthuriformes[27]<br>(–) Atheriniformes[10,12]<br>(–) Centrarchiformes[37]<br>(–) Cypriniformes[14]<br>(–) Cyprinodontiformes[15]<br>(–) Perciformes[16,17,18]<br>(ns) Perciformes[36†] | (–) Decapoda[30,35] | (–) Cardiida[41]<br>(–) Cardiida[32, 33]<br>(–) Venerida[34]<br>(ns) Venerida[55]<br>(ns) Pectinida[55]<br>(ns) Mytilida[54] | (–) Scleractinia[38] |
| | Clutch number | 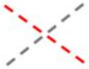 | 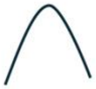 | (–) 11 Decapod spp.[43]<br>(–) 272 Rodent spp.[47‡]<br>(–) Marine invertebrates[52] | (+) Clupeiformes[13†] | | (–) Mytilida[54] | |
| | Clutch size | 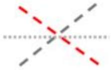 | 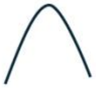 | (+) 1458 avian spp.[49‡]<br>(–) 11 Decapod spp.[43]<br>(ns) Sygnathiformes spp.[44]<br>(ns) Balanomorpha spp.[45] | (+) Acanthuriformes[27]<br>(+) Clupeiformes[13†]<br>(+) Perciformes[17,18]<br>(–) Perciformes[16]<br>(–) Atheriniformes[10,11]<br>(*) Centrarchiformes[37] | (+) Decapoda[3,4,5,6,8,28,30]<br>(–) Decapoda[7]<br>(+) Canuelloida[1,2]<br>(ns) Decapoda[9]<br>(*) Decapoda[39] | (+) Chitonida[26]<br>(+) Neogastropoda[22]<br>(+) Trochida[40] | (*) Actiniaria[31]<br>(ns) Echinoida[25]<br>(ns) Scleractinia[38] |
| **Reproductive output** | Fecundity | 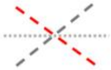 | 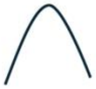 | (–) Marine invertebrates[52]<br>(ns) Scleractinia spp.[53] | (+) Atheriniformes[10,11]<br>(+) Perciformes[19]<br>(+) Salmoniformes[20†,21†]<br>(–) Clupeiformes[13†] | | | |
| **Offspring performance** | Offspring size | 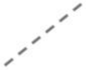 | 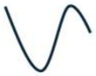 | (+) Sygnathiformes spp.[44]<br>(+) Marine/freshwater spp.[50] | | (+) Decapoda[7] | | |
| | Offspring provisioning | 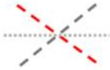 | 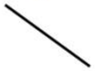 | (+) Sygnathiformes spp.[44]<br>(+) 278 fish spp.[46]<br>(+) 39 arthropod spp.[51]<br>(+) Marine invertebrates[52]<br>(ns/+) 10 teleost families[48]<br>(ns) Scleractinia spp.[53] | (+) Perciformes[17,23]<br>(–) Salmoniformes[20†,21†]<br>(ns) Atheriniformes[10] | (+) Isopoda[24]<br>(+) Decapoda[5,6,7,8,9, 28]<br>(–) Decapoda[3]<br>(ns) Decapoda[3, 39] | | |

[1]Lonsdale and Levinton 1985, [2]Lonsdale and Levinton 1986, [3]Baldzani et al. 2018, [4]Lardies and Wehrtmann 1997, [5]Lardies and Wehrtmann 2001, [6]Lardies and Castilla 2001, [7]Lardies et al. 2010, [8]Gorney et al. 1992, [9]Brante et al. 2004, [10]Conover 1992, [11]Sosebee 1991, [12]Middaugh and Hemmer 1992, [13]Leggett and Carscadden 1978, [14]Cowell and Resico 1975, [15]Conover and Present 1990, [16]Slesinger et al. 2021, [17]Kokita 2003, [18]Kokita 2004, [19]Richardson et al. 1997, [20]Fleming and Gross 1990, [21]Beacham and Murray 1993, [22]Waite et al. 2024, [23]Johnston and Leggett 2002, [24]Clarke and Gore 1992, [25]Lester et al. 2007, [26]Alvarez-Garcia et al. 2024, [27]Zarco-Perello et al. 2022, [28]Bezerra Ribeiro et al. 2023, [29]Domínguez-Petit et al. 2022, [30]Defeo and Cardoso 2002, [31]Ryan and Miller 2019, [32]Verdelhos et al. 2011, [33]Mahony et al. 2020, [34]Livore et al. 2019, [35]Bauer and Rivera Vega 1992, [36]Ishikawa and Kitano 2020, [37]Stocks et al. 2015, [38]de Putron and Smith 2011, [39]Stanski et al. 2018, [40]Martone and Micheli 2012, [41]Patiño et al. 2021, [42]Vila-Gispert et al. 2002, [43]Bauer 1992, [44]Foster and Vincent 2004, [45]Barnes and Barnes 1968, [46]Kasimatis and Ringos 2016, [47]Heldstab 2021, [48]Thresher 1988, [49]Boyer et al. 2010, [50]Marshall et al. 2018, [51]Thatje and Hall 2016, [52]Thorson 1950, [53]Gutierrez-Isaza et al. 2022, [54]Oyarzún et al. 2018, [55]Uribe et al 2012

Table S2. *Urosalpinx cinerea* field site information, specimen collector, number of specimens collected, date that breeding pairs were introduced to experimental conditions. Breeding pairs were established within one week of collection. NOAA Station IDs are the closest available sites used to gather publicly available data for calculating 3-year average daily regional temperatures.

| **Temperature Region** | **Collection site** | **U.S. State** | **Date paired** | **Latitude °N** | **Longitude °W** | **NOAA Station ID** |
|---|---|---|---|---|---|---|
| South | Skidaway | Georgia | 3/2/23 | 31.990 | 81.021 | 8670870 |
| South | Folly Beach | South Carolina | 3/2/23 | 32.660 | 79.943 | 8665530 |
| Mid | Beaufort | North Carolina | 3/2/23 | 34.718 | 76.671 | 8656483 |
| Mid | Cape Henlopen | Delaware | 4/6/23 | 38.789 | 75.102 | * |
| North | Fort Wetherill State Park | Rhode Island | 3/24/23 | 41.478 | 71.360 | 8452660 |
| North | Cape Cod | Massachusetts | 6/9/23 | 41.542 | 70.617 | 8447930 |
| North | Great Bay | New Hampshire | 6/9/23 | 43.107 | 70.863 | 8419870 |
| Cali | Richardson Bay | California | 4/13/23 | 37.894 | 122.497 | 9414290 |
| Cali | Coyote Point | California | 4/13/23 | 37.591 | 122.324 | 9414523 |
| Cali | Tomales Bay | California | 4/20/23 | 38.118 | 122.867 | 9415020 |
| Cali | Willapa Bay | Washington | 6/7/23 | 46.501 | 124.030 | * |

* NOAA sea surface temperature from OPS-API not used to calculate average regional temperature.

Table S3. Generalized linear mixed models implemented for each response variable using the “glmmTMB” R package, including zero-inflation and dispersion parameters to reduce effects of heteroscedasticity.

| **Response variable** | **glmmTMB distribution** | **zero-inflation[+] (ziformula)** | **dispersion parameter[†] (dispformula)** | **full model** |
|---|---|---|---|---|
| Total number of embryos | nbinom1 | ~1 | ~1 | ~ Population * Female size |
| Total number of capsules | nbinom1 | ~1 | ~1 | ~ Population * Female size |
| Total number of hatchlings | nbinom1 | ~1 | ~1 | ~ Population * Female size |
| Total number of clutches | nbinom1 | ~1 | ~1 | ~ Population * Female size |
| Offspring provisioning | lognormal(link = "log") | ~0 | ~Population | ~ Population + Female size + (1\| Female ID/clutch ID) |
| Offspring size | lognormal(link = "log") | ~0 | ~Population | ~ Population + Female size + (1\| Female ID/clutch ID) |
| Capsule size | gamma(link = “log”) | ~0 | ~Population | ~ Population + Female size + (1\| Female ID/clutch ID) |
| Embryo density | genpois(link = "log") | ~0 | ~Population | ~ Population + Female size + (1\| Female ID/clutch ID) |
| Offspring survival (F1) | betabinomial | ~1 | ~Population | ~ Population + Female size + (1\| Female ID/clutch ID) [‡] |
| Offspring survival (F2) | betabinomial | ~Population | ~1 | ~ Temperature * Population + Parental temperature + (1\| Female ID) [‡] |

[+]ziformula = ~0 (no zero-inflation), ~1 (probability equal for all observation), or ~factor; [†]dispformula = ~1 (standard dispersion for a given distribution family); [‡]weights = total number of embryos.
